# The contribution of Robos to olfactory sensory axon targeting in the zebrafish olfactory bulb

**DOI:** 10.64898/2026.03.19.709608

**Authors:** Jessica B. Herr, Emily S. Devereaux, Matthew J. Curran, Carly D. Seligman, Ryan P. Cheng, Jonathan A. Raper

## Abstract

Olfactory sensory neurons (OSNs) project a single axon from the olfactory epithelium to the olfactory bulb. OSNs initially target large, distinct, individually identifiable neuropils called protoglomeruli in the zebrafish embryo. Here we examine the contributions Robo axonal guidance receptors make to OSN axon targeting of protoglomeruli. We show that OSNs that project to the DZ protoglomerulus express higher levels of *robo2* than those that project to the CZ protoglomerulus, and concordant with this observation, DZ-projecting axons are more often misrouted by loss of *robo2* than are CZ-projecting axons. Further, we demonstrate that in the absence of *robo2*, *robo1* contributes to DZ-targeting but not to CZ-targeting. The loss of either *robo1* or *robo3* by themselves do not affect targeting to either the CZ or DZ protoglomeruli. These findings identify OSN subtype-dependent contributions of Robo receptors to vertebrate olfactory circuit assembly.

## Introduction

Understanding how neural circuit connectivity is established during development is central to deciphering how circuits function, and this is particularly conspicuous in the developing olfactory system. The first elements in the olfactory circuit, Olfactory Sensory Neurons (OSNs), originate in the olfactory epithelium. They project a single axon via the olfactory nerve to the olfactory bulb. Each OSN stochastically chooses to mono-allelically express approximately one Odorant Receptor (OR) from a large OR gene repertoire. In the zebrafish embryo, OSN axons initially target large, distinct, individually identifiable neuropils called protoglomeruli [1–2]. These well-defined targets facilitate genetic studies of olfactory axon guidance and targeting. This, coupled with the accessibility of all developmental stages and the optically transparent nature of the embryos [1,3], makes zebrafish an invaluable vertebrate system for studying the development of olfactory circuitry. Subsequent to the initial targeting of OSN axons to protoglomeruli, the axons of OSNs expressing the same OR coalesce together into distinct and reproducibly positioned neuropil regions known as glomeruli [4–6]. This developmental process forms a topographic map that transforms odorant experience into ordered patterns of odorant-specific neural activity in the olfactory bulb [7–9]. In this study, we systematically investigate the contribution that an important family of axon guidance receptors, the Robos, play in the formation of early olfactory circuitry in the larval zebrafish.

The initial protoglomerular target is tightly coordinated with OR choice. OSNs can choose to express main olfactory bulb type ORs from three OR-homology clades (A, B, and C) categorized by sequence similarity [10,11]. OSNs expressing ORs from clades A or B target a ventromedial region of neuropil in the olfactory bulb, the Central Zone (CZ) protoglomerulus. OSNs expressing ORs from clade C target a more dorsal and lateral neuropil, the Dorsal Zone (DZ) protoglomerulus. Single cell RNA-sequencing studies from our lab have identified genes that are differentially expressed between OSNs expressing ORs from different clades, suggesting coordination between the expression of axon guidance genes and OR choice [12,13].

In this study we determined the relative contributions that the axon guidance receptors *robo1*, *robo2*, and *robo3* play in CZ and DZ targeting. Robos are evolutionarily conserved single-pass transmembrane receptors that are important in many axonal guidance decisions [14–16]. Robo receptors generally induce chemorepulsive responses and are often activated by a family of highly conserved secreted Slit glycoproteins. In mammals there are three Slit genes (*slit1, slit2,* and *slit3*) and four Robo genes (*robo1*, *robo2*, *robo3*, and *robo4*). Robo-Slit signaling is required for the targeting of olfactory sensory axons in mouse [17] and in Drosophila [18]. Miyasaka et al found that in the zebrafish, *robo2* is essential for normal OSN axon pathfinding and reported that ubiquitous overexpression of *slit2* phenocopies loss of *robo2* [19]. Protoglomerular and glomerular targeting were shown to be grossly disrupted in these early studies. We previously identified *robo2* as being more highly expressed in OR clade C expressing, DZ projecting OSNs as compared to OR clade A- or B expressing, CZ projecting OSNs. We also reported that DZ, but not CZ projections are disrupted in *robo2* mutants [12,13]. Improved labelling of individual OSN axon trajectories and better clearing techniques now make a more detailed and accurate analysis of axon targeting defects in the absence of Robo and Slit activities possible.

Although *robo1* and *robo3* are also expressed in OSNs, their contributions to early OSN axon targeting are unknown. To explore the roles all the Robos play in protoglomerular targeting, we systematically knocked down each Robo or Slit using a CRISPR-based F0 knockdown approach. We assessed the accuracy of axon targeting in OR clade A expressing and CZ projecting OSNs, and in OR clade C expressing and DZ projecting OSNs. We found *robo2* to be more highly expressed in DZ than CZ projecting OSNs. Loss of *robo2* altered DZ projecting OSN axon trajectories more frequently than CZ axon trajectories. Loss of *robo1*, *robo3, slit2*, or *slit3* by themselves, or *slit2/slit3* together, did not induce axon targeting errors in either CZ or DZ projecting OSNs. Given the functional overlap among Slit ligands and their widespread expression in and around the olfactory nerve, the absence of *slit2*-dependent or *slit3*-dependent targeting defects suggests that other Slit family members may be sufficient to regulate OSN axon guidance. We show through genetic interaction experiments that *robo1* and *robo2* likely act together to guide DZ, but not CZ projecting axons to their targets. *Robo3* was found not to be required for accurate CZ or DZ projections. Together, our findings are consistent with a functional dose-dependent model in which Robo receptor expression levels tune OSN responsiveness to all Slits during early OSN targeting. We did not detect any specialization of Slit or Robo functions independent of expression level. These results suggest that Slits and Robos act redundantly to provide a simple mechanism by which diverse axon populations navigate a shared guidance landscape to locate specific targets.

## Materials and Methods

### Transgenic Zebrafish lines

Zebrafish lines were raised and maintained following the guidelines and under the supervision of the University of Pennsylvania Institutional Animal Care and Use Committee (IACUC, protocol #804895). University Laboratory Animal Resources (ULAR) at the University of Pennsylvania oversaw zebrafish veterinary care. Zebrafish larvae were raised at 28.5 °C and staged by hours post fertilization (hpf). The Yoshihara lab at the RIKEN Brain Science Institute generously provided the Tg(omp:lyn-RFP)^rw035a^ and Tg(trpc2:gap-Venus)^rw036a^ zebrafish lines [19,20]. The Tg(UAS:gap43-citrine)^p201Tg^ line was described by Lakhina et al [21]. The Tg(BACOR111-7:IRES:GAL4)^zf1062Tg^ and Tg(BACOR130-1:IRES:GAL4)^zf1063Tg^ lines were described by Shao et al [11].

The robo2^ti272z^ (*astray*) mutant was described in Fricke et al and genotyped using a KASP assay (Biosearch Technologies) (KASP sequence: 5’ - GGTTATAGCACTGGCCTCTGTGTGTCTCTCTTTACAGATATAAGTCCtCCAGCGCAGGGTGTGGATCAC[A/T]GACATGTGCAaAAGGAGCTGGGAgAAGTTATTGTGCGGCTACaCAACCCTGTGGTCCTTAGCCCTACCACCATACAGGTCACATGGACGGTAAGACGCTC - 3’) [22]. The nell2a^sa32506^ and nell2b^sa18051^ mutants were obtained from The Zebrafish International Resource Center (ZIRC). Embryo heads of the same genotype were pooled and processed for image acquisition.

### F0 Knockdown Experiments

Three CRISPR-Cas9 RNP complexes targeting different sites in the gene were used to generate first-generation knockdowns [23]. CRISPR guide-RNAs (crRNAs) were selected that cut sequences in functionally important exons in the first half of the targeted gene. CHOPCHOP was used for crRNA design, and cutting efficiency and indel production were predicted using CRISPRscan [24,25]. Four candidate CRISPR targets were screened via headloop PCR for each target gene [23]. The actual efficiency of selected guides was quantified using direct sequencing data subjected to indel analysis by DECODR [26].

Embryos were generated by crossing Tg(BacOR111-7:IRES:Gal4) or Tg(BACOR130- 1:IRES:Gal4) fish with Tg(UAS:gap43-citrine) fish. Their progeny were injected with a mixture of 3 validated RNP complexes at the single-cell stage. All embryos were maintained at 28.5°C until 72hpf, at which point they were anesthetized, euthanized, and prepared for imaging.

The sequences of each crRNA, headloop PCR primers, and DECODR sequencing primers are provided in **Supplemental File 1**.

### Image Acquisition

Larvae were fixed overnight at 4°C in 4% paraformaldehyde (PFA) dissolved in 0.1 M phosphate buffer. Subsequently, a CUBIC-based tissue clearing protocol was performed [27]. Larvae were immersed and gently shaken in CUBIC 1 solution for 2 hours at room temperature. Following 3 five-minute washes in PBS and 1 ten-minute wash in PBS-Triton x-100, the treated larvae were nuclear-stained with RedDot2 in PBS (1:75 by volume; Biotium 40061) overnight. The larvae were washed for 5 minutes 3 times in PBS and immersed in CUBIC 2 solution for at least 2 hours, mounted, and imaged in CUBIC 2 solution.

Images were obtained using a Leica SP5 confocal microscope equipped with a 63x oil-immersion objective. Z-stacks were generated with a step size of 1μm beginning at the anterior margin of the olfactory epithelium and extending to the posterior limit of the olfactory bulb. Processing and visualization of the resulting z-stacks was performed with FIJI/ImageJ and ClearVolume [28,29].

### Quantification of axonal targeting errors

Imaging data were anonymized and protoglomerular targeting was scored blind by two independent observers. Discrepancies in scoring were resolved by the scorers prior to unblinding. Axon termination sites were scored relative to their expected protoglomerular target. Ectopic misprojections were quantified independently across the anterior-posterior, dorsal-ventral, and medial-lateral axes. Each error was classified as Branching, Escape, or Direct. A branching error consists of an axon with a single identifiable branch point and 2 or more OSN branches terminating in different locations. Axons with an escape error project to their expected protoglomerular target but have a termination outside of the protoglomerular border. Axons displaying a direct error have a single axon that projects to any location outside of the expected protoglomerular target. There is no overlap between these categories. The statistical significance of misprojections was assessed via a two-tailed Fisher’s exact test.

### Single label *In situ* hybridization

*In situ* probe templates for *slit1a* (Addgene plasmid # 61279; http://n2t.net/addgene:61279; RRID:Addgene_61279) and *slit1b* (Addgene plasmid # 61280 ; http://n2t.net/addgene:61280; RRID:Addgene_61280) were a generous gift from Chi–Bin Chien & Lara Hutson [30]. *In situ* probe templates for *slit2* (5’_tcaccaagatcccagaccac_3’ and 5’_gcagctcggacatgttactg_3’) and *slit3* (5’_aaggcttgttcgctcctctg_3’ and 5’_atcaaacgcaccctcacgga_3’) were generated by PCR amplification from 48hpf zebrafish cDNA. The amplicons were each cloned into a pCR4-TOPO vector using a TOPO TA Cloning Kit (Invitrogen, 45-0030). The probes were generated using a DIG RNA labeling kit (Sigma-Aldrich, 11175025910) and subsequently purified by LiCl/Ethanol precipitation.

Embryos were generated by crossing Tüpfel Long Fin (TLF; ZDB-GENO-990623-2) fish together for 48hpf slit preparations. They were collected, raised at 28.5°C, anesthetized, euthanized, and then fixed overnight at 4°C in 4% PFA in 0.1 M phosphate buffer. They were then dehydrated in a methanol series and stored in methanol at -20°C.

Embryos were rehydrated by 2 five-minute washes in DEPC-PBST. Prior to *in situ* hybridization, zebrafish larvae were treated with proteinase K (10 uL of 1mg/mL in 1 mL DEPC- PBST) for 23 minutes at 25°C. Following digestion, embryos were washed for 5 minutes in DEPC-PBST, fixed in 4% PFA dissolved in DEPC-PBS for 20 minutes, and then washed 2 times in DEPC-PBST for 5 minutes. DEPC-PBST was replaced with DEPC-H2O and quickly replaced with a mixture of 2.5uL of acetic anhydride in 1mL of 0.1M triethanolamine and incubated for 10 minutes at room temperature. Embryos were then washed 2 times for 10 minutes each in DEPC-PBST. Each slit expression pattern was detected using digoxigenin (DIG) labelled RNA antisense probes hybridized overnight at 57°C. After hybridization, embryos were soaked 2 times for 30 minutes each in 50% formamide in 2X SSCT at 57°C, rinsed for 15 minutes in 2XSSCT, and rinsed 2 times for 30 minutes each in 0.2X SSCT. Embryos were then incubated in 0.3% H2O2 in PBS at room temperature for 30 minutes to inactivate endogenous peroxidases and then washed 5 times for 5 minutes each in PBST at room temperature. Following blocking in 10% Fetal Bovine Serum in PBST for 3-4 hours at room temperature, the embryos were incubated in anti-DIG-POD (1:500 in 10% Fetal Bovine Serum in PBST; Roche, 11207733910) at 4°C overnight. Following 4 25-minute PBST washes at room temperature, the label was amplified utilizing the cyanine 5-coupled tyramide kit (1:50; Akoya Biosciences, NEL745001KT) according to instructions. In the 36hpf embryos, citrine-expressing axons were visualized with goat anti-GFP (1:100; Rockland Immunochemicals, 600–101–215) followed by donkey anti-goat IgG Alexa Fluor 488 (1:500; Invitrogen) as previously described by Lakhina et al (2012). All preparations were then washed 2 times for 5 minutes each in 2X SSC and then incubated in propidium iodide dissolved in 2X SSC for 8 minutes at room temperature (330ug/ml, Sigma-Aldrich, P4170) to stain nuclei. They were then washed 6 times for 3 minutes each in 2X SSC, fixed for 20 minutes in 4% PFA dissolved in PBS, and washed 2 times for 5 minutes each in PBST. Finally, animals were mounted for imaging in Vectashield mounting medium (Vector Laboratories, H-1000) and imaged using a Leica SP5 Confocal Microscope. All z-stacks were acquired using identical acquisition settings for each probe.

### Dual Label *In Situ* Hybridization

Robo and OR probe sequences were approximately 900 bp long and were embedded between KpnI/EcoRI in a pUC18 vector as follows: *robo1* (RefSeq accession number NM_001365167) nucleotides 473-1372, *robo2* (RefSeq accession number NM_131633) nucleotides 1244-2143, *robo3* (RefSeq accession number NM_001328416) nucleotides 485-1384, *OR130-1* (RefSeq accession number NM_001110317) nucleotides 132-1031, and *OR111-7* (RefSeq accession number NM_131582) nucleotides 226-1125.

The dual label *in situ* hybridization protocol matches the single label *in situ* hybridization protocol until just before probe hybridization. Each OR expression pattern was detected using Fluorescein labelled RNA antisense probes and each robo expression pattern was detected using digoxigenin (DIG) labelled RNA antisense probes. The probes were hybridized overnight at 57°C. After hybridization, embryos were soaked 2 times for 30 minutes each in 50% formamide in 2X SSCT at 57°C, rinsed for 15 minutes in 2XSSCT, and rinsed 2 times for 30 minutes each in 0.2X SSCT. Embryos were then incubated in 0.3% H2O2 in PBS at room temperature for 30 minutes to inactivate endogenous peroxidases and then washed 5 times for 5 minutes each in PBST at room temperature. Following blocking in 10% Fetal Bovine Serum in PBST for 3-4 hours at room temperature, the embryos were incubated in anti-fluorescein-POD (1:500 in 10% Fetal Bovine Serum in PBST; Roche, 11426346910) at 4°C overnight. Following 4 25-minute PBST washes at room temperature, the label was amplified utilizing a fluorescein-coupled tyramide kit (PerkinElmer, NEL741001KT) according to instructions. Embryos were then washed 3 times for 5 minutes each at room temperature in PBST, followed by a quick rinse in PBS for 1 minute, an incubation in 1% H2O2 in PBS at room temperature for 30 minutes, and 5 washes for 5 minutes each in PBST. Following blocking in 10% Fetal Bovine Serum in PBST for 3-4 hours at room temperature, the embryos were incubated in anti-DIG-POD (1:500 in 10% Fetal Bovine Serum in PBST; Roche, 11207733910) at 4°C overnight. Following 4 25-minute PBST washes at room temperature, the label was amplified utilizing the cyanine 5-coupled tyramide kit (1:50; Akoya Biosciences, NEL745001KT) according to instructions. The embryos were then washed 2 times for 5 minutes each in 2X SSC and then incubated in propidium iodide dissolved in 2X SSC for 8 minutes at room temperature (330ug/ml, Sigma-Aldrich, P4170) to stain nuclei. They were then washed 6 times for 3 minutes in 2X SSC, fixed for 20 minutes in 4% PFA dissolved in PBS, and washed 2 times for 5 minutes each in PBST. Finally, animals were mounted for imaging in Vectashield mounting medium (Vector Laboratories, H-1000) and imaged using a Leica SP5 Confocal Microscope. All z-stacks were acquired using identical acquisition settings.

Neurons labelled with a single OR probe are sparse within the olfactory epithelium. Individual labelled OSNs were first identified and outlined in FIJI/ImageJ. Z-sections spanning the whole cell were flattened by addition before measuring the mean robo fluorescence intensity for each outlined OSN. The mean background fluorescence was measured in the whole olfactory epithelium from the same flattened image. The background intensity was subtracted from the OSN measurement to obtain the final background-corrected mean fluorescence value. The statistical significance of mean fluorescence between groups was assessed via a two-tailed Mann Whitney test.

## Results

### *Robo2* F0 knockdown phenocopies the *robo2* mutant *astray*

We used a CRISPR-based F0 knockdown approach to elucidate the roles of Robo receptors and their Slit ligands in OSN axon pathfinding. This approach has been demonstrated to phenocopy a wide range of null mutations affecting early development [23]. The axons targeting specific protoglomeruli were visualized using two transgenic lines that label distinct subsets of OSNs, TRPC2-Venus and OMP:RFP. Ciliated olfactory sensory neurons express Main Olfactory Bulb type ORs and Olfactory Marker Protein b (OMPb), while microvillous sensory neurons express V2R-type odorant receptors and TRPC2 [20]. Each class terminates in distinct and mutually exclusive, individually identifiable protoglomeruli (**Fig 1A**) [20]. OMP expressing OSNs project axons to the Central Zone (CZ), Dorsal Zone (DZ), Lateral Glomerulus 3 (LG3), and Medial Glomerulus (MG). TRPC2 is expressed in OSNs whose axons project to the Olfactory Plexus (OP), Lateral Glomerulus 1, 2, and 4 (LG1, LG2, LG4), and the Ventral Posterior Glomerulus (VPG). The mutant zebrafish line *astray^ti272z^* (*ast*) lacks a functional *robo2* receptor [22]. It was crossed into transgenic lines containing both OMP:RFP and TRPC-Venus. OSN axon trajectories in *ast* homozygous mutant embryos have fully penetrant and severe axon guidance defects in both OMP- and TRPC2-labelled OSNs (**Fig 1B**). Many labelled OSN axons reach the olfactory bulb but the resulting protoglomeruli appear disorganized. A subset of OSN axons misroute posteriorly and in most embryos some of them cross the midline to the contralateral olfactory bulb (**Fig 1B cyan arrow**). Midline-crossing errors were never observed in wild-type embryos. These observations confirm previous findings that *robo2* is required for proper OSN axon guidance and protoglomerular targeting in zebrafish [12,19].

**Fig 1.**
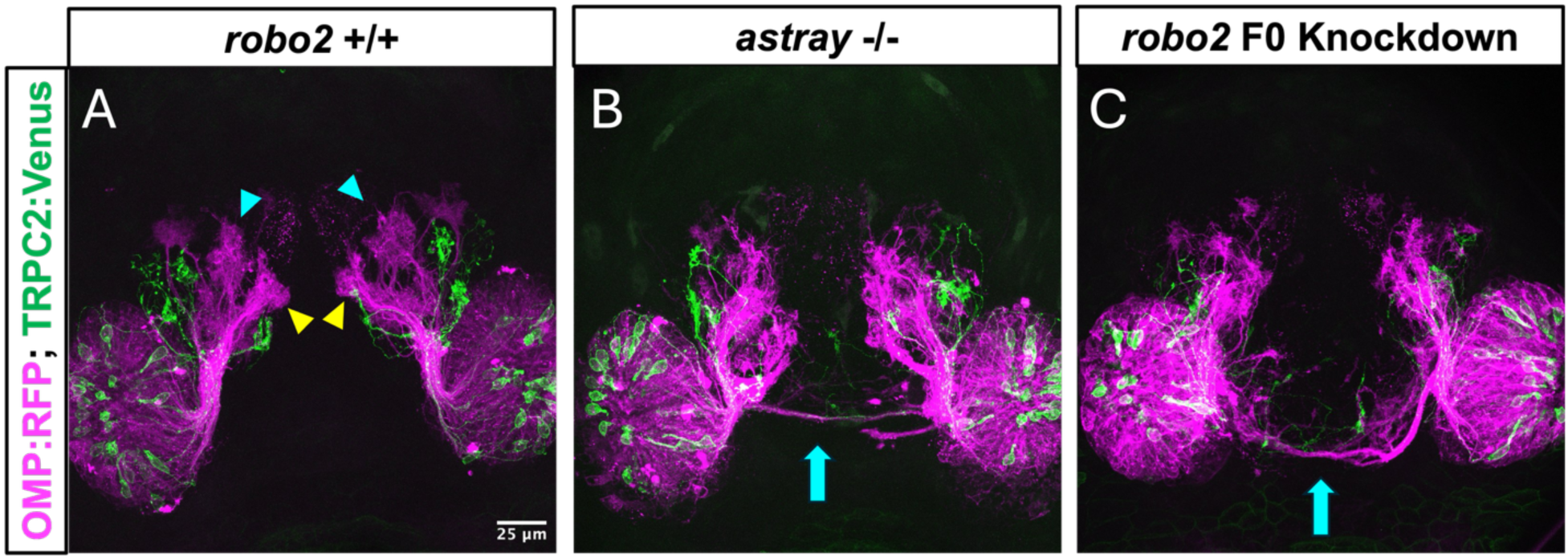
*Robo2* F0 knockdown qualitatively phenocopies *astray*. **(A)** Representative projection from an optical z-stack of the zebrafish wild-type olfactory system at 72 hpf. Specific non-overlapping subsets of OSNs (see text) are labeled by OMP:RFP (Magenta) and TRPC2:Venus (Green). OMP-expressing axons project to the Central Zone (CZ, indicated by yellow arrowheads), Dorsal Zone (DZ, indicated by cyan arrowheads), Lateral Glomerulus 3 (LG3), and Medial Glomerulus (MG). TRPC2-expressing axons project to the Olfactory Plexus (OP), Lateral Glomerulus 1, 2, and 4 (LG1, LG2, LG4), and the Ventral Posterior Glomerulus (VPG). **(B)** Protoglomeruli appear disorganized and a subset of labelled OSN axons misroute posteriorly in *ast* homozygous mutants and **(C)** *robo2* F0 knockdowns. Midline crossing to the contralateral side is observed in *ast* homozygous mutants or *robo2* F0 knockdowns as indicated by cyan arrows.

To determine whether a multi-CRISPR *robo2* F0 knockdown reliably phenocopies the *ast* mutant, we injected three RNPs targeting *robo2* into OMP:RFP; TRPC2:Venus transgenic embryos at the one-cell stage. Assuming they act independently, the three RNPs targeting *robo2* should together introduce indels with frameshifts or premature stops in 96% of targeted *robo2* genes **(Table 1)**. F0 knockdown of *robo2* induces severe axonal targeting defects in both OMP- and TRPC2-labelled OSNs similar to the defects observed in *ast* mutants, (**Fig 1C**). The strength and penetrance of the *robo2* knockdown phenotype was also similar to that of the *ast* homozygous mutant. These results suggest that the CRISPR-based F0 knockdown approach reliably reduces *robo2* function in a manner comparable to the *ast* homozygous mutant.

**Table 1:**
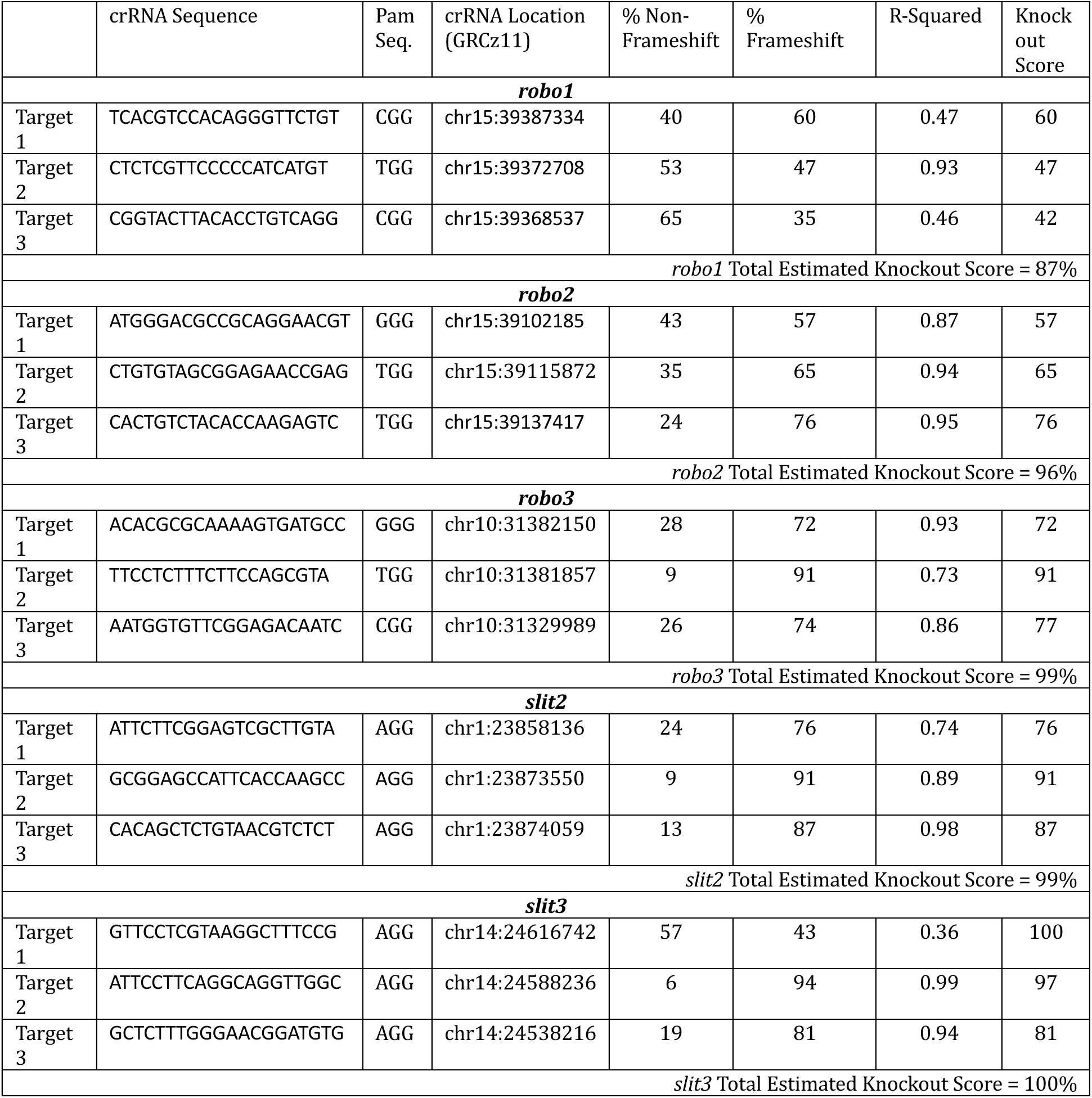
CRISPR F0 knockdown effectiveness as determined by DECODR. For each Robo or Slit gene the effectiveness of the top three crRNA targets was assessed by DECODR in pools of 10 embryos at 48hpf. The knockout score represents the frequency of frameshift mutations or induced premature stops. R-squared represents the goodness-of-fit between the predicted edit composition and the observed Sanger sequencing trace generated by DECODR.

### *Robo2* contributes to the protoglomerular targeting of specific subsets of OSNs that target different protoglomeruli

A zebrafish line containing Tg(BACOR111-7:IRES:Gal4) (subsequently referred to as BacOR111-7) crossed with a line containing the fluorescent reporter Tg(UAS:gap43-Citrine) (subsequently referred to as UAS:Citrine) label a small number of OR111-7 clade A expressing OSNs whose axons project to the ventral and medially located CZ protoglomerulus (**Fig 2A**). Tg(BACOR130-1:IRES:Gal4) (subsequently referred to as BacOR130-1) crossed with the UAS:Citrine line label a small number of OR130-1 clade C expressing OSNs whose axons project to the dorso-laterally located DZ protoglomerulus (**Fig 2C**). These transgenic lines enable us to assess the accuracy of OSN axon trajectories to CZ and DZ protoglomeruli.

**Fig 2.**
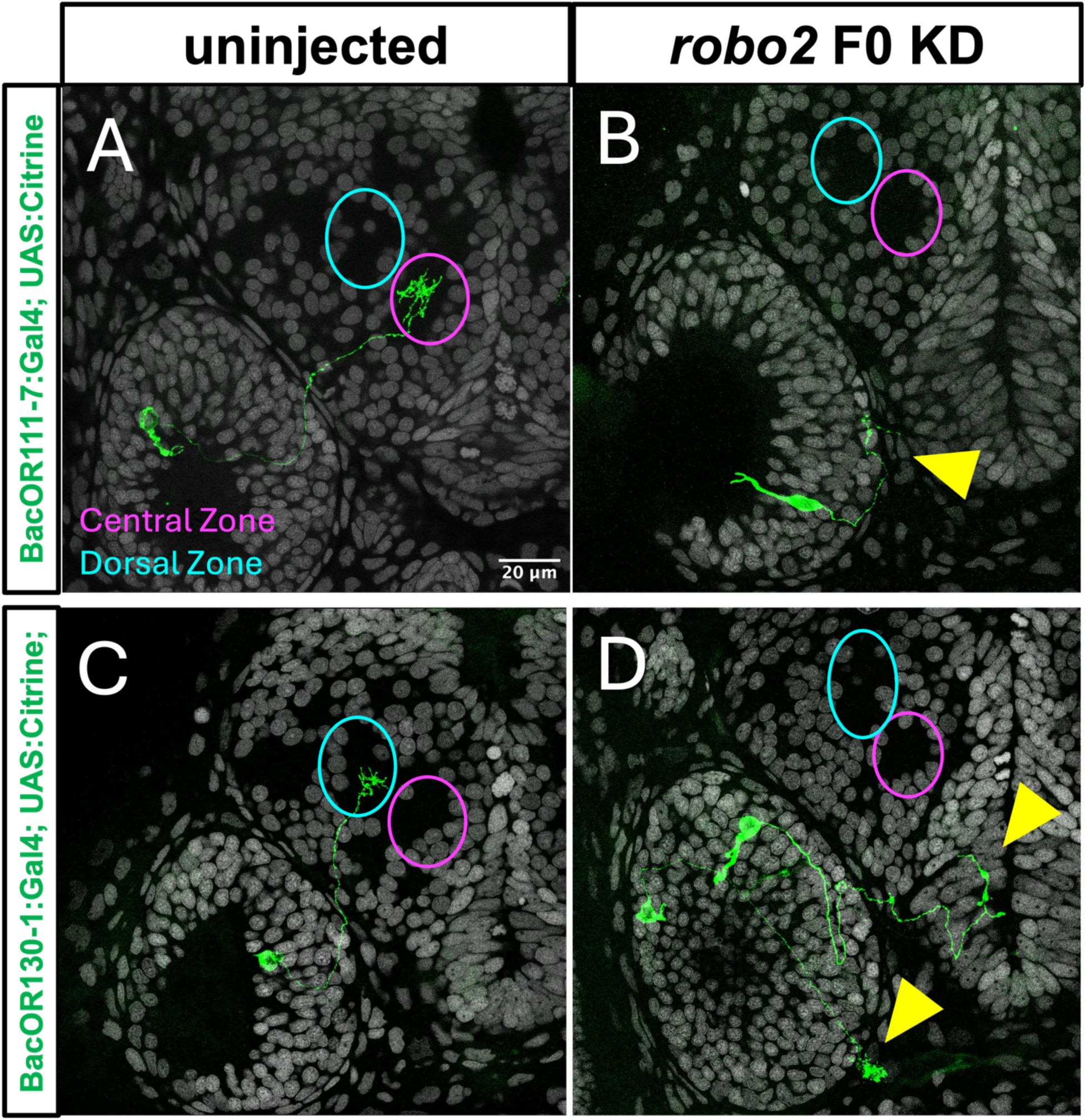
F0 knockdown of *robo2* induces ventral and posterior errors in both DZ and CZ projecting OSNs. BacOR111-1 CZ-projecting or BacOR130-1 DZ-projecting OSN axon trajectories were visualized at 72hpf in either uninjected embryos, or sibling embryos injected with RNPs that knockdown *robo2*. **(A)** The axons of BacOR111-7 labeled OSNs terminate in the CZ protoglomerulus (magenta) in uninjected embryos but **(B)** frequently terminate ventrally and posteriorly in *robo2* F0 knockdowns. **(C)** The axons of BacOR130-1 labeled OSNs terminate in the DZ protoglomerulus (cyan) in uninjected embryos but **(D)** frequently terminate ventrally and posteriorly in *robo2* F0 knockdowns. Yellow arrowheads indicate misprojecting axons.

To determine the contribution of *robo2* to OSN axon targeting, *robo2* was knocked down at the one-cell stage and labelled axons were visualized in both CZ-projecting BacOR111-7 and DZ- projecting BacOR130-1 OSNs. F0 knockdown of *robo2* resulted in both CZ and DZ projection errors (**Fig 2B, 2D**). The predominant errors were direct misprojections to ectopic ventral and posterior locations (**Supplemental Fig 1**). Misprojecting axons also occasionally crossed the midline. Other misprojections consisted of ectopic terminations within the olfactory epithelium.

### Quantitative analysis of *robo1*, *robo2*, or *robo3* knockdowns in CZ or DZ projecting OSNs

To quantitatively test if *robo1* or *robo2* are required for the correct targeting of subsets of OSN axons, knockdown experiments were performed in both DZ-projecting BacOR130-1 and CZ- projecting BacOR111-7 labelled embryos. Assuming they act independently, the three RNPs targeting *robo1* should together introduce indels with frameshifts or premature stops in 87% of *robo1* genes **(Table 1)**. F0 knockdown of *robo1* did not affect the percentages of labelled OSN axons making errors in either DZ-projecting BacOR130-1 or CZ-projecting BacOR111-7 axons as compared to uninjected sibling controls. In contrast, F0 knockdown of *robo2* significantly increased the percentage of labelled OSN axons making errors for both DZ-projecting BacOR130-1 and CZ-projecting BacOR111-7 OSNs (**Fig 3A, 3B**).

**Fig 3.**
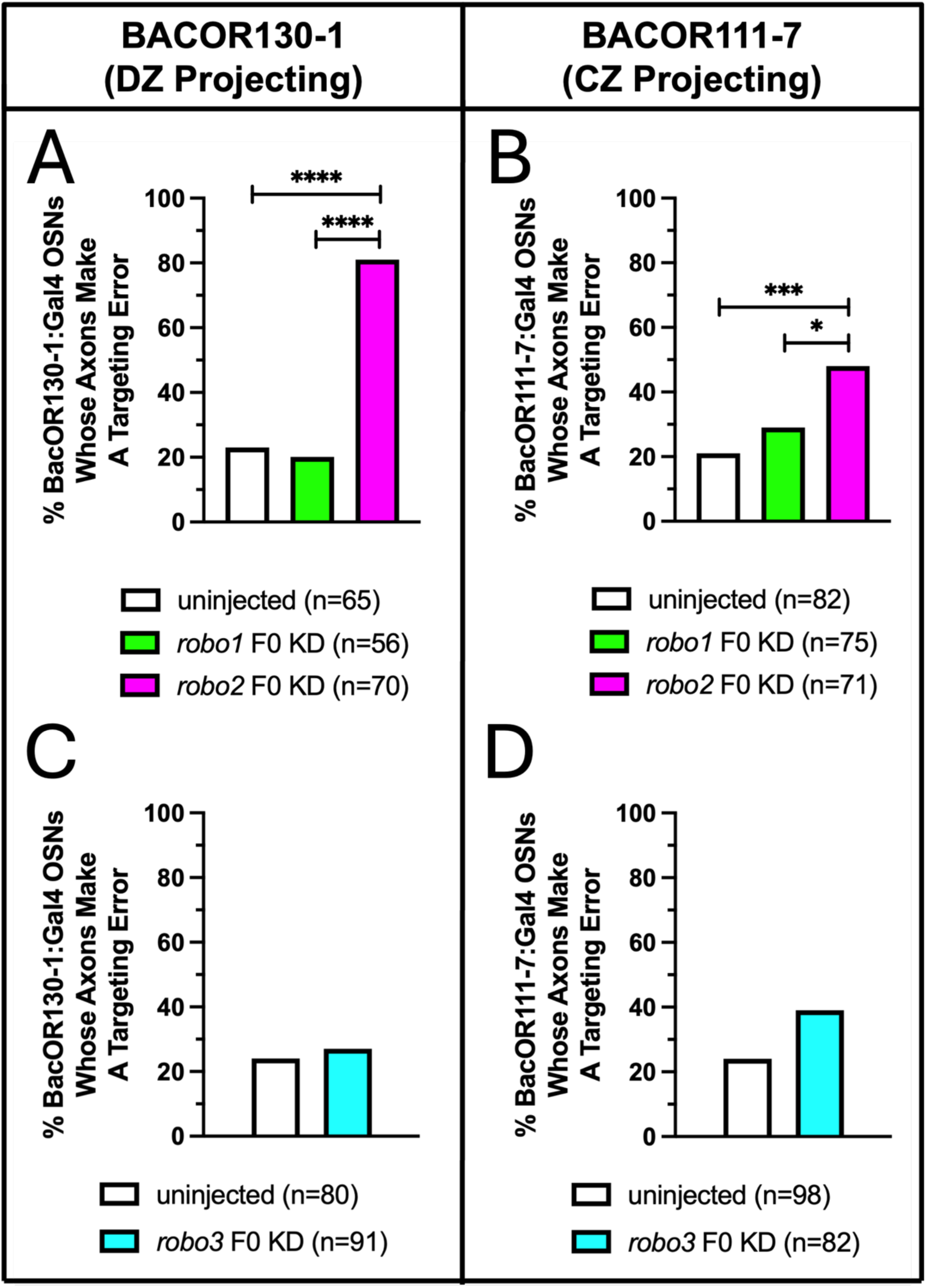
F0 knockdown of *robo2* disrupts axonal projections in OSNs. OSN axon trajectories of uninjected embryos and sibling embryos injected with RNPs that knockdown *robo1*, *robo2*, or *robo3* were compared at 72hpf. **(A, B)** There is no statistical difference in the percentage of DZ-projecting BacOR130-1 or CZ-projecting BacOR111-7 OSNs whose axons make targeting errors in *robo1* F0 knockdown embryos as compared to uninjected sibling controls. F0 knockdown of *robo2* significantly increased the percentage of labelled axons making an error in both DZ-projecting BacOR130-1 and CZ-projecting BacOR111-7 OSNs. **(C, D)** There is no statistical difference in the percentage of DZ-projecting BacOR130-1 or CZ-projecting BacOR111-7 OSNs whose axons make targeting errors in *robo3* F0 knockdown embryos as compared to uninjected sibling controls. Statistical comparisons were performed using a two- tailed Fisher’s Exact test with * denoting p ≤ 0.05, *** denoting p ≤ 0.001, and **** denoting p ≤ 0.0001.

*Robo3* F0 knockdowns were performed in both DZ-projecting BacOR130-1 and CZ-projecting BacOR111-7 OSNs. Assuming they act independently, the three RNPs targeting *robo3* should together introduce indels with frame shifts or premature stops in 99% of *robo3* genes **(Table 1)**. F0 knockdown of *robo3* did not affect the percentages of OSNs making errors in DZ-projecting BacOR130-1 or CZ-projecting BacOR111-7 axons compared to uninjected sibling controls (**Fig 3C, 3D**).

### *Robo2* is more highly expressed in DZ as compared to CZ projecting OSNs

We performed dual-label fluorescent *in situ* hybridization (FISH) to determine whether higher expression of *robo2* in OR clade C as compared to OR clade A and B expressing neurons reflects expression patterns in OR 130-1 and OR 111-7 OSNs [12,13]. We compared the relative expression levels of *robo1*, *robo2*, or *robo3* in the cell bodies of CZ projecting OR111-7, or DZ projecting OR130-1 expressing OSNs at 48hpf when these axons are actively projecting into protoglomeruli. OSNs labeled with a probe to a single OR are sparsely distributed in the olfactory epithelium. We identified individual OSNs labelled with either a probe for OR111-7 or OR130-1. The density of the fluorescent signal for Robo expression was then quantified for the selected cell and compared to the average signal in a larger area of the epithelium **(Fig 4A**, see Methods for quantification). Neither *robo1* nor *robo3* expression levels are measurably different between OR111-7 or OR130-1 expressing OSNs in two separate experiments (**Fig 4B, 4D**). In contrast, *robo2* is significantly more highly expressed in OR130-1 expressing, DZ projecting OSNs as compared to OR111-7 expressing, CZ projecting OSNs **(Fig 4C)**. These data are consistent with RNAseq data where *robo2* was more highly expressed in OSNs expressing clade C ORs as compared to clade A ORs [12,13].

**Fig 4.**
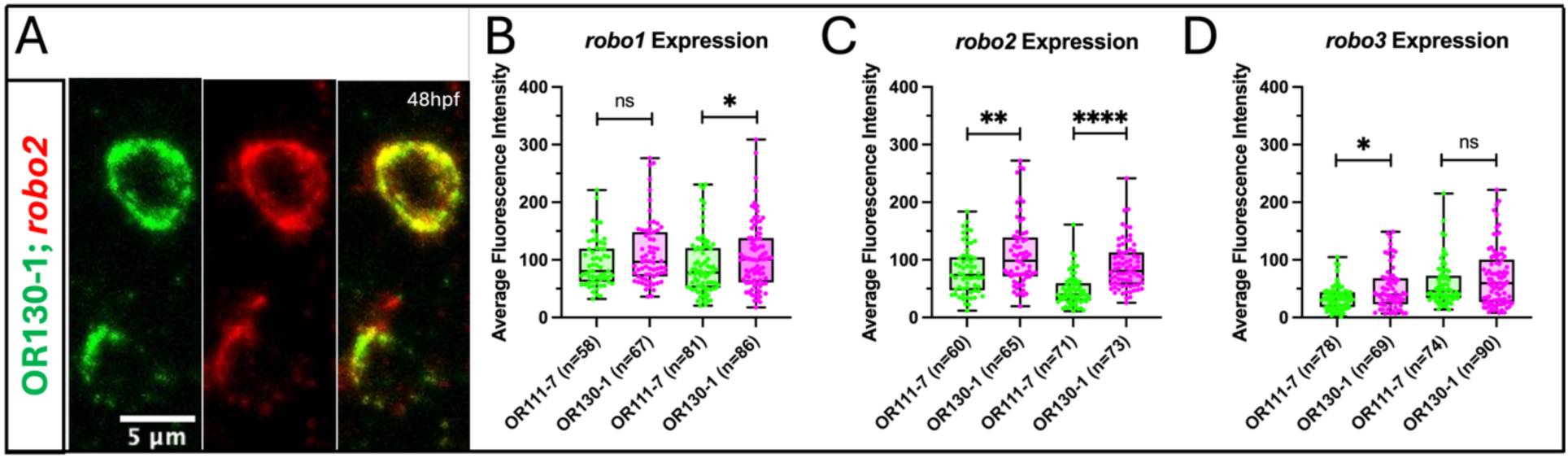
Differential *robo* expression in OSNs assessed by dual fluorescent *in situ* hybridization. **(A)** Dual-FISH of *robo2* co-expressed with an OR130-1 probe. Average fluorescence intensities in OR130-1 or OR111-7 OSNs is reported for *robo1* **(B)**, *robo2* **(C)**, or *robo3* **(D)**.

### *Robo2* F0 knockdown quantitatively phenocopies *astray*

To further validate the F0 knockdown approach, we sought to determine whether the *robo2* F0 knockdown quantitatively phenocopies the *ast* mutant. Fish containing *ast* +/-; BacOR130-1 were crossed to fish containing *ast* +/-; UAS:Citrine. Half of the resulting just-fertilized embryos were injected with the three *robo2* F0 RNPs and the other half were used as uninjected sibling controls. Citrine labelled OSN axon trajectories were compared for four different conditions: (1) uninjected *ast* +/+, (2) uninjected *ast* -/-, (3) *robo2* F0 KD injected *ast* +/+, and (4) *robo2* F0 KD injected *ast* -/-. The percentage of protoglomerular targeting errors in uninjected *ast* -/- embryos was not significantly different from either *ast* +/+ embryos injected with *robo2* F0 KD or *ast* -/- embryos injected with *robo2* F0 KD **(Fig 5A)**. We conclude that *robo2* F0 KD has a quantitatively similar phenotype as compared to the *ast* mutant. That *robo2* F0 KD in *ast* mutant embryos has a similar phenotype as compared to *ast* mutants suggests that both the *robo2* F0 KD and *ast* phenotypes represent a full loss of *robo2* function.

**Fig 5.**
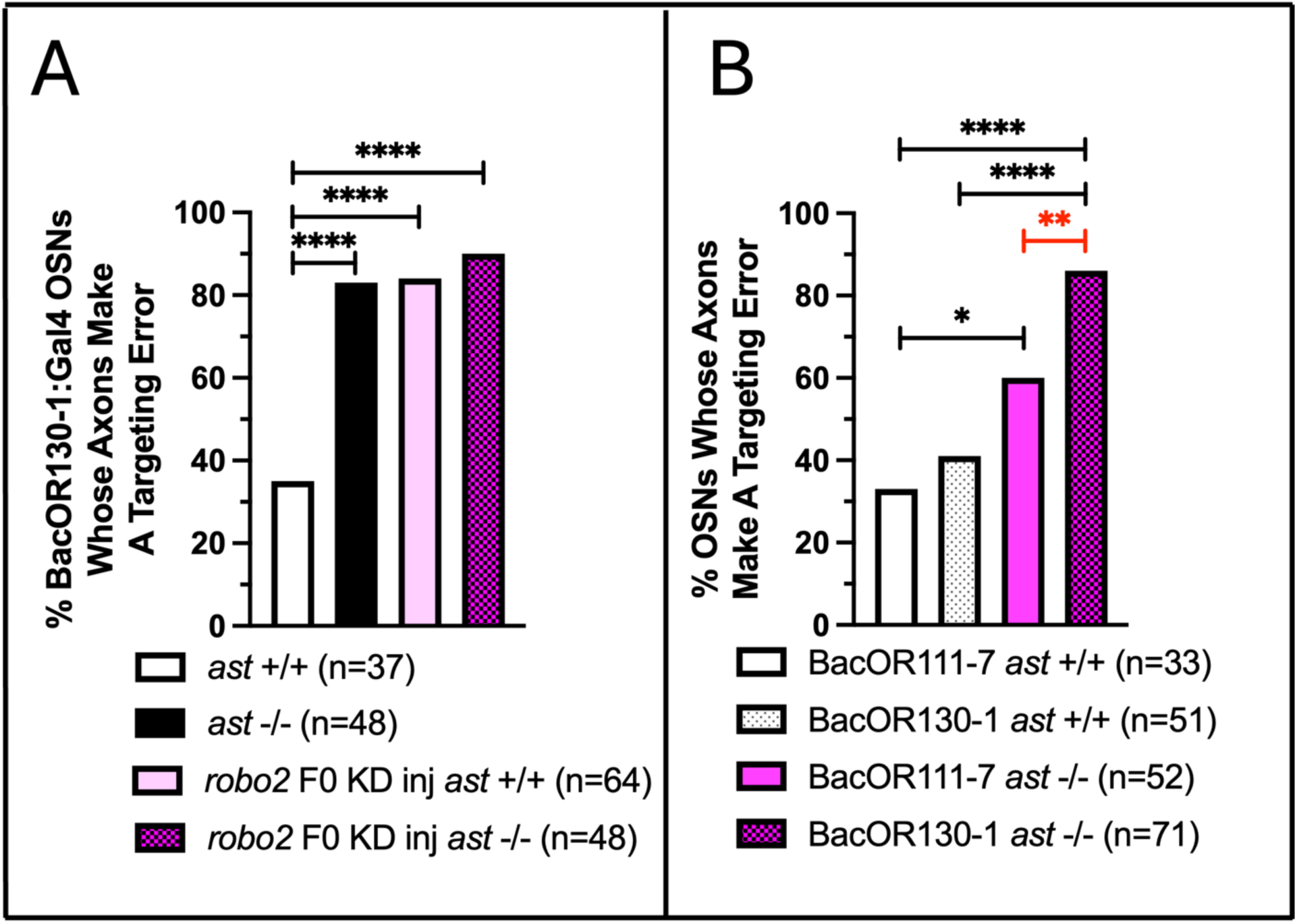
*Robo2* quantitatively phenocopies *ast* and loss of *robo2* has a more severe effect on DZ-projecting BacOR130-1 than CZ-projecting BacOR111-7 OSN axons. (A) The percentages of DZ-projecting BacOR130-1 OSN axons making targeting errors in *ast* +/+ or *ast* -/- embryos with or without *robo2* F0 knockdown as compared to sibling uninjected embryos. **(B)** The percentages of DZ-projecting BacOR130-1 or CZ-projecting BacOR111-7 expressing OSNs whose axons exhibit targeting errors in *ast* +/+ or *ast* −/− embryos. Statistical comparisons were performed using a two-tailed Fisher’s Exact test with * denoting p ≤ 0.05, ** denoting p ≤ 0.01, and **** denoting p ≤ 0.0001.

### *Robo2*’s contributions to DZ as compared to CZ OSN pathfinding

The dual-label FISH shows that *robo2* is significantly expressed at higher levels in OR130-1 expressing, DZ projecting OSNs as compared to OR111-7 expressing, CZ projecting OSNs. *Robo2* F0 knockdown resulted in an increased error rate in DZ as compared to CZ axon projections **(Fig 3A, 3B)**. To further explore the possibility that DZ projecting OSN axons are more dependent on *robo2* function than CZ projecting OSN axons, fish containing *ast* +/-; BacOR130-1; BacOR111-7 were crossed to fish containing *ast* +/-; UAS:Citrine. The citrine labelled axon trajectories were compared in the resulting progeny at 72hpf for four different conditions: (1) *ast* +/+; BacOR130-1, (2) *ast* +/+; BacOR111-7, (3) *ast* -/-; BacOR130-1, and (4) *ast* -/-; BacOR111-7. The percentages of labelled DZ-projecting BacOR130-1 and CZ- projecting BacOR111-7 OSNs making targeting errors was not significantly different between *ast* +/+ embryos **(Fig 5B)**. In contrast, DZ-projecting BacOR130-1 OSN axons made significantly more errors than CZ-projecting BacOR111-7 OSN axons in *ast* -/- embryos. This further supports the conclusion that DZ projections are more sensitive to a loss of *robo2* than CZ projections.

### Knockdown of *robo1* in *astray* mutants

We next sought to determine if the combined loss of *robo2* and *robo1* produces additive, synergistic, or redundant effects on OSN axon targeting. Fish containing either *ast* +/-; BacOR111-7, or *ast* +/-; BacOR130-1, were crossed to fish containing *ast* +/-; UAS:Citrine. Half of the resulting just-fertilized embryos were injected with *robo1* F0 KD RNPs. The remaining embryos were used as uninjected sibling controls. This design resulted in six experimental conditions. Citrine labelled axon trajectories and protoglomerular targeting in *ast* +/+, +/-, and -/- progeny were compared in *robo1* F0 knockdown or uninjected controls. There is a significant increase in BacOR130-1 axons mistargeting the DZ protoglomerulus in *robo1* F0 KD; *ast* -/- as compared to uninjected *ast* -/- embryos **(Fig 6A)**. In contrast, there is no observed increase in BacOR111-7 labelled axons mistargeting the CZ protoglomerulus in *robo1* F0 KD; *ast* -/- as compared to uninjected *ast* -/- embryos **(Fig 6B)**. Similar to those in *ast* -/- embryos, the errors in BacOR130-1 *robo1* KD ast -/- embryos were direct errors to ventral and posterior locations **(Supplemental Fig 2)**. These data indicate that *robo1* can contribute to BacOR130-1 DZ targeting in a manner similar and redundant to *robo2*. *Robo1* makes no detectable contribution to CZ targeting in BacOR111-7 OSNs even in the absence of *robo2*.

**Fig 6.**
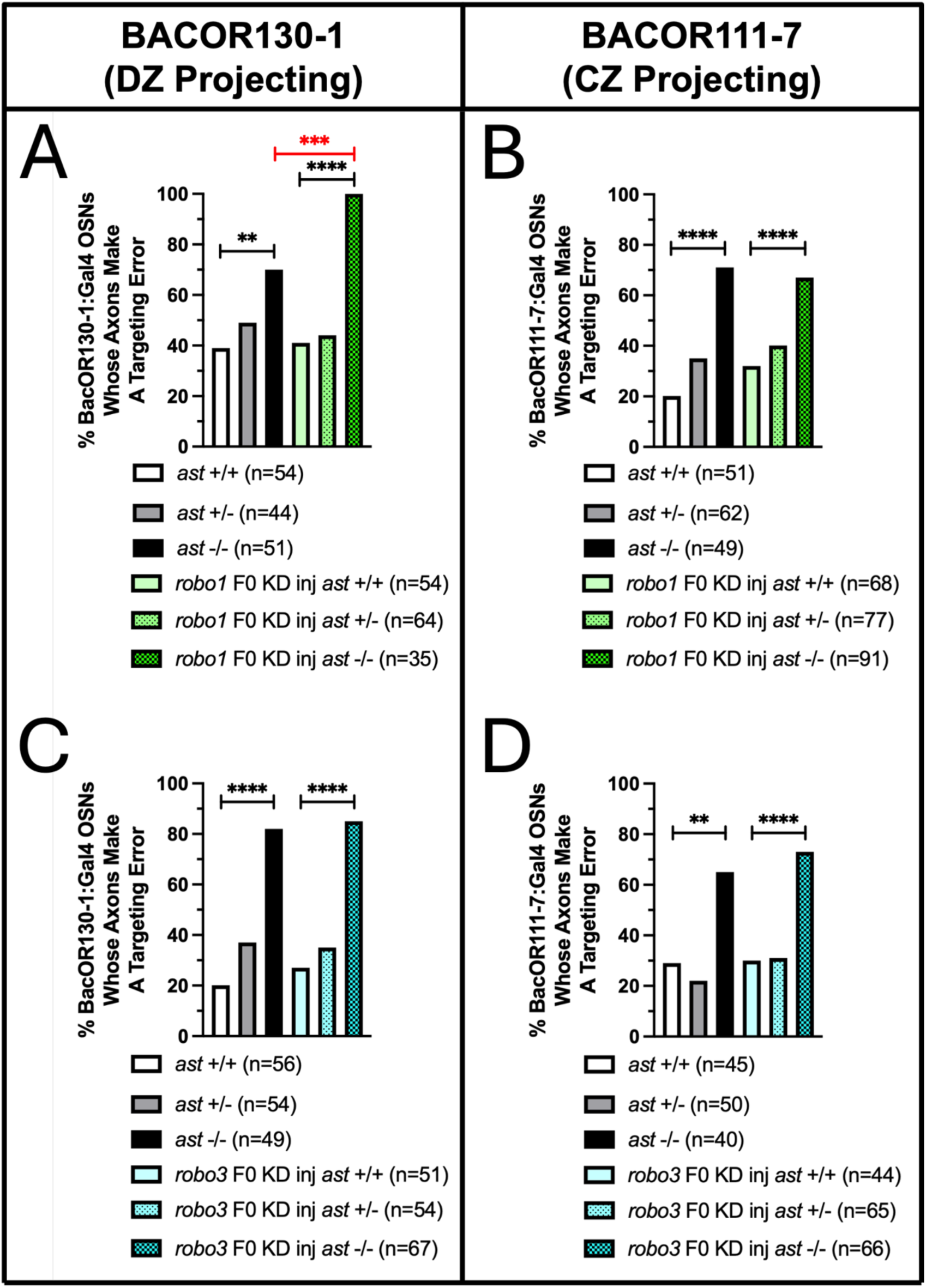
*Robo1* provides partial redundancy to *robo2* in BacOR130-1 but not in Bac111-7 OSNs. The percentages of axon targeting errors in *ast* +/+, +/-, or -/- embryos in *robo1* F0 knockdowns or uninjected controls in **(A)** DZ-projecting BacOR130-1 or **(B)** CZ-projecting BacOR111-7 OSNs. The percentages of axon targeting errors in *ast* +/+, +/-, or -/- embryos in *robo3* F0 knockdown or uninjected controls in **(C)** DZ-projecting BacOR130-1 or **(D)** CZ- projecting BacOR111-7 OSNs. Statistical comparisons were performed using a two-tailed Fisher’s Exact test with * denoting p ≤ 0.05, ** denoting p ≤ 0.01, *** denoting p ≤ 0.001, and **** denoting p ≤ 0.0001.

### Knockdown of *robo3* in *astray* mutants

We next sought to determine if the combined loss of *robo2* and *robo3* produces additive, synergistic, or redundant effects on OSN targeting. Fish containing either *ast* +/-; BacOR111-7 or *ast* +/-; BacOR130-1 were crossed to fish containing *ast* +/-; UAS:Citrine. Half of the resulting just-fertilized embryos were injected with *robo3* F0 KD RNPs. The remaining embryos were used as uninjected sibling controls. This design resulted in six experimental conditions. Citrine labelled axon trajectories and protoglomerular targeting in *ast* +/+, +/-, and -/- progeny were compared in both uninjected embryos and embryos subjected to *robo3* F0 knockdown. There is no observed increase in protoglomerular mistargeting of DZ- or CZ-projecting axons in *robo3* F0 KD; *ast* -/- as compared to uninjected *ast* -/- embryos **(Fig 6C, 6D)**.

### Slits are expressed in distinct patterns within the olfactory bulb

*In Situ* Hybridization (ISH) was performed to visualize slit expression patterns at 48hpf in wholemount embryos as previously described [21]. Each image in Figure 7 is a representative 3-µm maximum intensity projection from an individual preparation (See Methods). All four slits are expressed within or near the zebrafish olfactory bulb at 48hpf. *Slit1a* is expressed most strongly in cells throughout the olfactory bulb **(Fig 7A)**. *Slit1b* is expressed in cells within the olfactory bulb and appears specifically enriched in cells near the CZ protoglomerulus **(Fig 7B** cyan stars). *Slit2* is most highly expressed ventrally near the midline **(Fig 7C)**. Finally, *slit3* is also most highly expressed ventrally near the midline but to a lesser extent than *slit2* **(Fig 7D)**.

**Figure 7.**
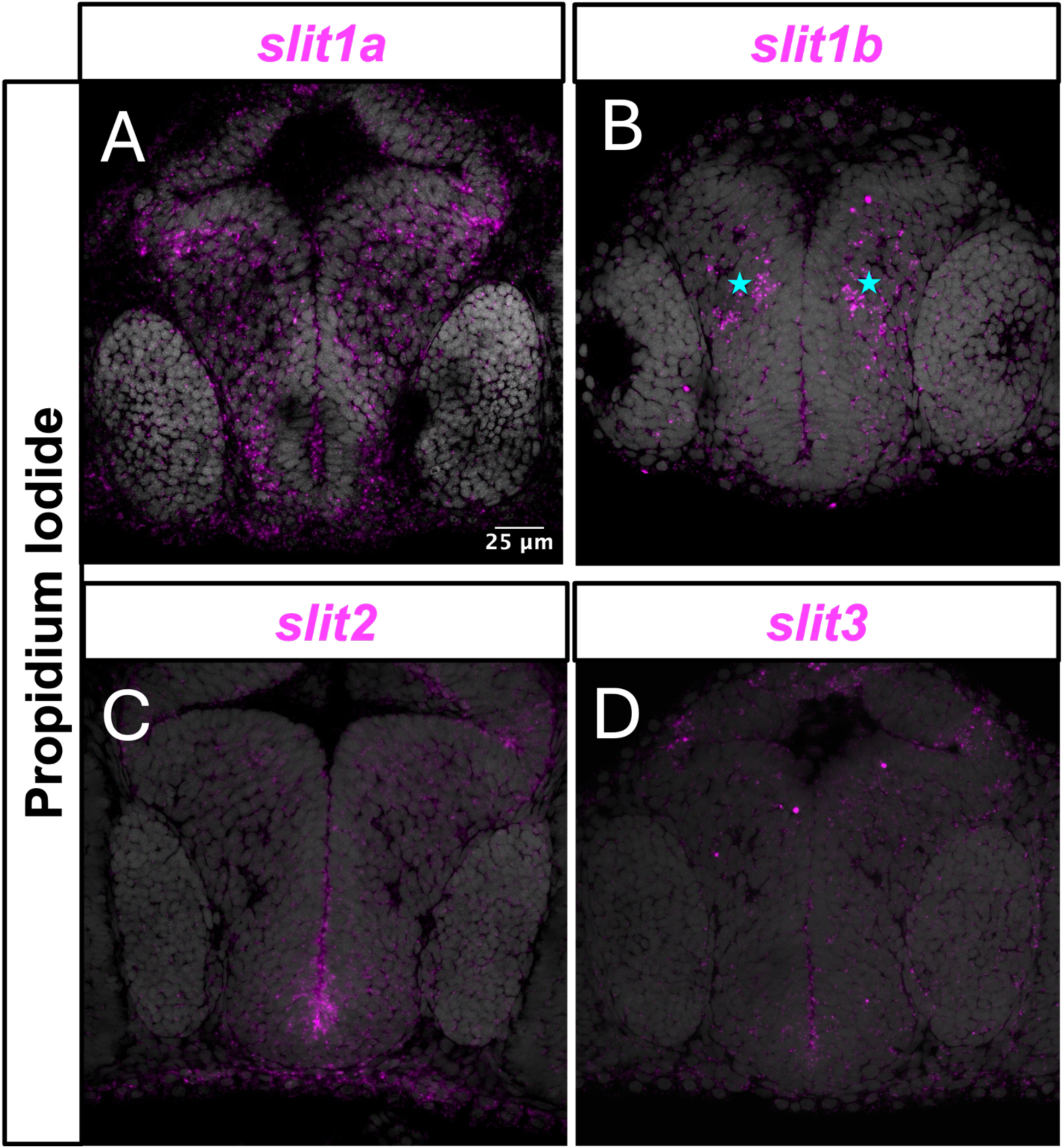
Expression patterns of *slit1a*, *slit1b*, *slit2*, and *slit3* in the embryonic olfactory bulb. Fluorescent *in situ* hybridization (Magenta) was performed in 48 hpf embryos. Representative maximum-intensity projections from 3-µm z-stacks in individual preparations are shown. Cyan stars indicate CZ protoglomeruli surrounded by *slit1b* expression.

### Knockdown of *slit1a*, *slit1b*, *slit2* , slit3, *slit2* and *slit3* together, or *slit1a* and *slit1b* together do not induce axon targeting errors

Both *slit2* and *slit3* are enriched along the midline in the ventral forebrain and are hypothesized to promote dorsally directed outgrowth and prevent midline crossing [19]. Ubiquitous overexpression of *slit2* in transgenic zebrafish impairs OSN pathfinding in a similar manner to that observed an *ast* homozygous mutant [19]. These findings led to the proposal that *slit2* is the likely ligand required for *robo2* activity in OSN guidance. We sought to determine if *slit2* is required for the correct targeting of OSN axons. Knockdown experiments were performed in both DZ-projecting BacOR130-1 and CZ-projecting BacOR111-7 OSNs. Assuming they act independently, the three RNPs targeting *slit2* and the three RNPS targeting *slit3* should introduce indels with frameshifts or premature stops in 99% of targeted genes **(Table 1)**. However, neither F0 knockdown of either *slit2* or *slit3* induce axon targeting errors in DZ projecting BacOR130-1 or CZ projecting BacOR111-7 OSNs (**Fig 8A, 8B**). We also performed *slit1a* and *slit1b* knockdown experiments in CZ-projecting BacOR111-7 OSNs. The three RNPs targeting *slit1a* and the three RNPs targeting *slit1b* were found to produce indels through a headloop PCR (Kroll et al., 2021). However, neither F0 knockdown of *slit1a* or *slit1b* induce axon targeting errors in CZ-projecting BacOR111-7 OSNs (**Fig 8C**).

**Fig 8.**
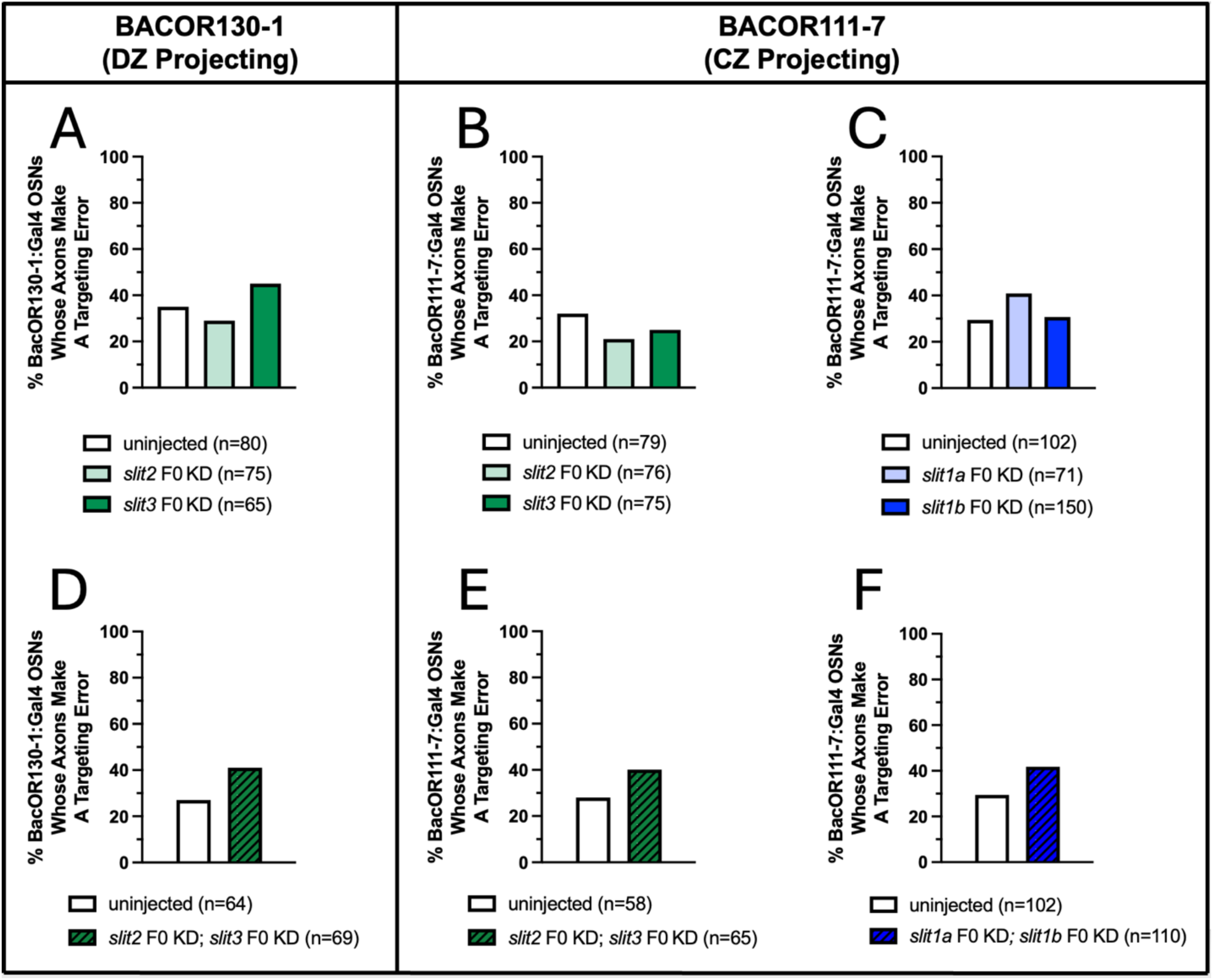
F0 knockdown of *slit1a*, *slit1b*, *slit2*, *slit3*, *slit1a/slit1b* together, or *slit2/slit3* together do not disrupt OSN targeting. The trajectories of OSNs in *slit1a, slit1b, slit2*, or *slit3* knockdown embryos were compared to uninjected sibling controls at 72hpf. **(A, B)** Neither *slit2* or *slit3* knockdown induced errors in DZ-projecting BacOR130-1 or CZ-projecting BacOR111-7 OSNs. **(C)** Neither *slit1a* or *slit1b* knockdown induced errors in CZ-projecting BacOR111-7 OSNs. **(D, E, F)** Similar results were obtained when both *slit2 and slit3* were knocked down together or when both *slit1a* and *slit1b* were knocked down together. Statistical comparisons were performed using a two-tailed Fisher’s Exact test.

Disruption of *slit3* expression in a *slit2 -/-* background causes severe defects comparable to those observed in *robo2* mutants at the zebrafish optic chiasm [31]. This led to the conclusion that zebrafish *slit2* and *slit3* act together to regulate retinal axon crossing at the midline. We completed *in situ* hybridization experiments in *slit2* KD embryos and observed a potential upregulation of *slit1a* and *slit3* upon loss of *slit2* **(Supplemental Fig 3)**. Since we did not observe a phenotype in either the *slit2* or *slit3* knockdowns and both are expressed at the ventral midline, we hypothesized that these two slit ligands may act together to regulate OSN axon targeting in the zebrafish olfactory bulb. A comparison of DZ-projecting BacOR130-1 or CZ-projecting BacOR111-7 axon trajectories in uninjected and *slit2*/*slit3* double knockdowns was performed to determine if their combined loss would produce a phenotype similar to knockdown of *robo2*. To avoid the toxicity that can occur when too many RNPs are injected into embryos, the two best gRNAs for *slit2* were combined with the two best gRNAs for *slit3* and injected into just fertilized embryos. We sequenced targeted regions from a pool of 10 embryos to determine the knockdown efficacy from each injected batch used to score axonal trajectories. The sequencing results indicated the knockdowns were 99% effective for each slit. There is no observed increase in DZ-projecting BacOR130-1 or CZ-projecting BacOR111-7 axon mistargeting compared to uninjected sibling controls **(Fig 8D, 8E)**. We also compared CZ- projecting BacOR111-7 axon trajectories in uninjected and *slit1a*/*slit1b* double knockdowns to determine if their combined loss would produce a phenotype similar to knockdown of *robo2*. Similar to the double knockdown of *slit2*/*slit3* together, we observed no increase in CZ- projecting BacOR111-7 axon mistargeting in *slit1a*/*slit1b* double knockdowns as compared to uninjected sibling controls **(Fig 8F)**. These results indicate that either additional non-slit ligands contribute to the *robo2* phenotype; or alternatively, the four slit paralogs may function redundantly and the loss of any single slit gene, or pair of genes, is compensated for by the remaining paralogs.

## Discussion

We have performed both positive and negative control experiments to validate the CRISPR based F0 knockdown approach used in this study. To assess whether F0 knockdown itself affects OSN targeting, we previously knocked down the pigmentation gene *slc24a5* (solute carrier family 24 member 5) as a negative control [13]. Mutations in *slc24a5* are associated with reduced pigmentation in zebrafish and its function is not expected to affect OSN axon guidance [32]. Despite the obvious depletion of pigmentation in the retina of F0 *slc24a5* knockdown embryos, both DZ-projecting BacOR130-1 and CZ-projecting BacOR111-7 OSN axons projected normally when compared to uninjected controls. This indicates the F0 knockdown approach does not in itself cause an increase in targeting errors and that uninjected embryos are suitable controls for experiments. As a positive control, we show here that knockdown of *robo2* both qualitatively and quantitatively phenocopies the *ast* mutant. This phenotypic concordance demonstrates that F0 knockdown can be both efficient and functionally equivalent to stable mutant lines. Our previous study showed that knockdown of *pcdh11* qualitatively and quantitatively phenocopies a *pcdh11* mutant [13]. These results support the use of F0 knockdown as a reliable and time-efficient method for dissecting complex axon guidance signaling systems. One limitation of the CRISPR based approach we employed is that it generates whole body knockouts. Interpretation of the results in this study is based on the reasonable assumption that Robos act cell autonomously in OSN cells, but we cannot rule out indirect non-autonomous effects from Robo-related influences on cell adhesion, identity, differentiation, or survival [33].

Robos have previously been shown to play a crucial role in the guidance of longitudinal axon tracts in *Drosophila*. The positioning of these tracts was hypothesized to be governed by a combinatorial Robo receptor code [34,35]. *Robo* is expressed in the most medial of the longitudinal axon tracts, *robo* and *robo3* are expressed in intermediate tracts; and *robo*, *robo2*, and *robo3* are expressed in the most lateral tracts. The loss of either *robo2* or *robo3* induces lateral tract displacement closer to the midline, whereas forced expression of *robo2* or *robo3* in medial axons displaces them away from the midline. These experiments led to the hypothesis that the positioning of longitudinal axonal trajectories are specified by distinct combinations of Robo receptors that confer graded sensitivity to Slit ligands expressed at the midline. Later work in *Drosophila* refined this model further through the analysis of Robo gene swaps [36]. This experimental strategy involved driving the expression of one Robo receptor in the pattern of another, directly testing whether Robo family members are functionally interchangeable. Interestingly, the lateral positioning of longitudinal axon tracts was largely normal in all of the Robo receptor swaps. These results suggested that axon guidance decisions are reliant on differences in Robo gene regulation and not dependent on distinct functional properties of individual Robo receptors.

Our findings are consistent with a similar model in which individual Robo receptors do not have unique specialized properties. Slit-Robo signaling is one of many contributing factors that affect the trajectories of olfactory sensory axon extension towards the olfactory bulb in zebrafish [19]. Sensory axons begin to emerge from the olfactory epithelium and grow towards the olfactory bulb from 24-36 hpf, a time that coincides with strong *robo2* mRNA expression in the olfactory placode. Pioneering OSN axons subsequently arrive at the olfactory bulb around 48 hpf and begin to form protoglomeruli as *robo2* expression in the olfactory epithelium begins diminishing. This expression pattern suggests *robo2* functions primarily during early OSN axon outgrowth and pathfinding. We found *robo2* to be required for axon targeting of both DZ-projecting BacOR130-1 and CZ-projecting BacOR111-7 OSNs. However, DZ-projecting BacOR130-1 OSNs express higher levels of *robo2* and are more severely affected by its loss than CZ- projecting BacOR111-7 OSNs.

Importantly, the loss of individual Robo receptors other than *robo2* does not induce changes in OSN axon trajectories, contrary to what might be predicted if individual Robos make unique or specialized contributions to OSN axon guidance. Loss of individual Robo receptors does not redirect OSN axons to distinct alternative locations. Loss of *robo1* produces no observable phenotype in CZ-projecting BacOR111-7 OSNs, and in these neurons, loss of *robo1* and *robo2* together simply phenocopy *robo2* single knockouts. In contrast, loss of *robo1* accentuates the *robo2* loss of function phenotype in DZ-projecting BacOR130-1 OSNs. As in *Drosophila*, our data are consistent with a functional dose-dependent model in which Robo receptors tune the magnitude of slit responsiveness during early OSN targeting, rather than encoding positional identity through distinct receptor functions. This framework provides a very simple mechanism by which diverse axon populations navigate a shared guidance landscape to achieve precise wiring.

Robo receptors guide mouse OSN axons by mediating repulsive responses to slit ligands expressed within the developing olfactory bulb. The canonical ligands for Robos are the Slits which in the mouse are expressed in the developing olfactory bulb and ventral forebrain. They have been shown to regulate OSN axon trajectory and targeting [37,38]. For example, early-arriving, dorsal-innervating, *robo2*-positive OSN axons navigate to the dorsal portion of the olfactory bulb via repulsive interactions with *slit1* expressed in the ventral olfactory bulb [17,39,40]. In zebrafish, previous *in situ* hybridization experiments revealed that all 4 slit paralogs are expressed in distinct patterns throughout the olfactory bulb with *slit2* and *slit3* specifically enriched at ventral and posterior midline locations [19]. Ubiquitous overexpression of *slit2* resulted in a phenotype nearly identical in strength and penetrance to the *ast* zebrafish mutant, consistent with the notion that early OSN pathfinding depends on the spatial distribution of Slits. We found that knockdown of *slit1a*, *slit1b*, *slit2*, *slit3*, *slit1b/slit1b* together, or *slit2*/*slit3* together all failed to induce axon targeting defects. These results suggest that either an unidentified Robo ligand plays a key role in OSN axon guidance, or that all slits function redundantly, so that the loss of any single slit gene is compensated for by the remaining paralogs. In this context, the upregulation of both *slit1a* and *slit3* upon the loss of *slit2* may be significant. An alternative Robo ligand, NELL2, can bind both *robo2* or *robo3* in the mouse [41–43]. However, we found no observable misprojection phenotype in OMP-expressing or TRPC2- expressing OSN axons in *nell2a*-/-;*nell2b*-/- fish at 72hpf compared to uninjected sibling controls **(Supplemental Fig 4)**. Since Robo activation by misexpressed Slit2 clearly affects OSN pathfinding, a reasonable conclusion from our results is that there is a profound redundancy between Slit ligands in determining OSN axon trajectories.

OSN axon misprojections in *robo2* mutants are biased toward ventral and posterior locations of the olfactory bulb. This suggests that Slit-Robo signaling repels OSN axons from these inappropriate regions. Its absence might allow these axons to be attracted into ectopic locations by another guidance system. We have shown that *netrin1a, netrin1b*, *sema3fa*, *sema3d*, *pcdh11*, *pcdh7b, pcdh10b, pcdh17*, and all the slit paralogs are expressed in the olfactory bulb during early OSN axon outgrowth and pathfinding [3,13,21,44]. Thus, early OSN axon growth cones are exposed to a mixed guidance environment containing a multitude of repulsive and attractive cues. Future genetic interaction experiments will be required to determine whether other guidance pathways are necessary or sufficient to drive the mistargeting observed in *robo2* mutants.

These results add to the growing body of evidence that many guidance cues are involved in precise protoglomerular targeting, as Slit–Robo signaling likely operates within a broader network of cell surface and ECM protein interactions. We have shown that many classical guidance factors play a role in early axon guidance in the developing olfactory system, such as Netrin/Deleted in Colorectal Cancer (DCC) [21,45], Slit/Robo [12,19], Semaphorin/Neuropilin [3,44], and non-clustered protocadherins [13]. An axon’s targeting is determined by the collective actions of many different independent guidance processes. Recent work in *Drosophila* olfactory circuit assembly demonstrated that the loss of individual guidance cues or receptors often have poorly penetrant effects, and that it is necessary to systematically alter combinations of sensory cell surface proteins to reprogram axon targeting [46,47]. These findings and our work support the hypothesis that proper and robust neural connectivity emerges from the additive activity of many weakly attractive and repulsive axon guidance cues acting together.

---

**Fig S1.**
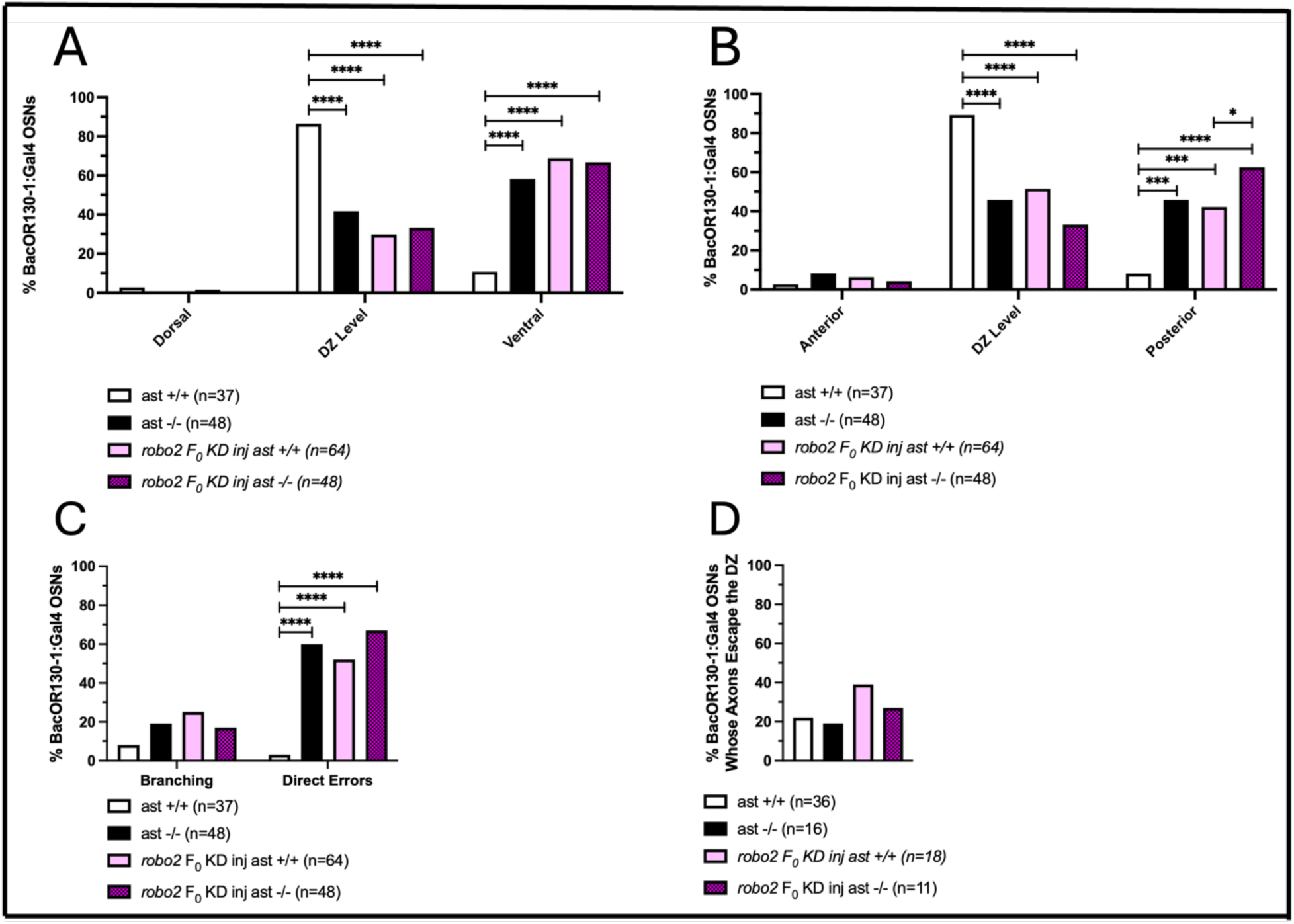
The predominant OSN axon errors in *robo2* F0 KD and *ast* were direct misprojections to ectopic ventral and posterior locations. (A, B) DZ-projecting BacOR130- 1 labeled axons in *robo2* F0 KD embryos and *ast*-/- were more likely to terminate in ectopic ventral or posterior locations in relation to the DZ protoglomerulus than in uninjected controls. **(C, D)** Misprojections of DZ-projecting BacOR130-1 labeled axons in *robo2* F0 KD embryos and *ast*-/- were more likely to be classified as direct errors instead of branching or escapes from the DZ protoglomerulus. Statistical comparisons were performed using a two-tailed Fisher’s Exact test with * denoting p ≤ 0.05, *** denoting p ≤ 0.001, and **** denoting p ≤ 0.0001.

**Fig S2.**
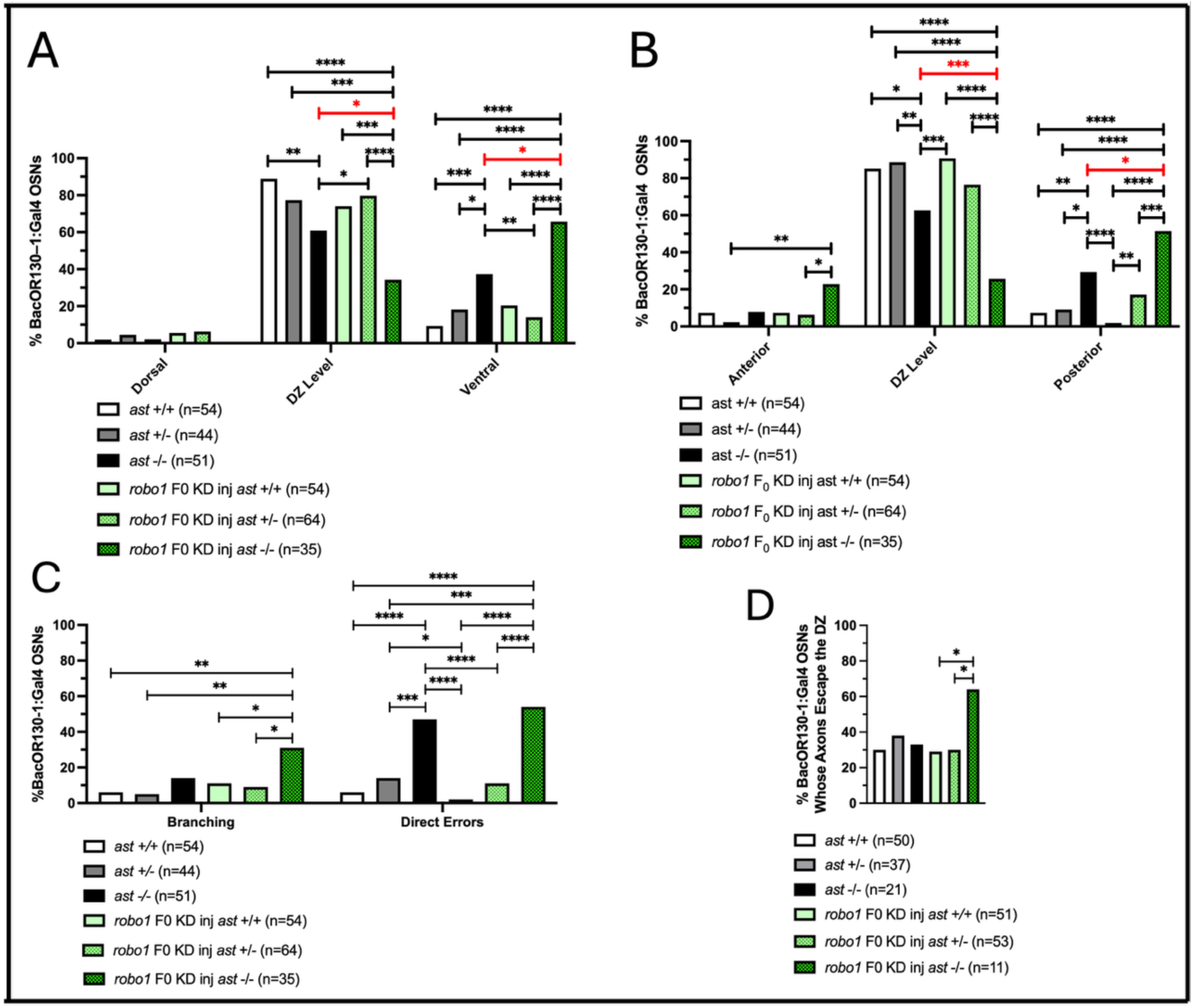
The predominant errors in *robo1* F0 KD; *ast*-/- embryos are direct misprojections to ectopic ventral and posterior locations in DZ-projecting BacOR130-1 OSNs. (A) DZ- projecting BacOR130-1 labeled axons in *robo1* F0 KD; *ast*-/- were more likely to terminate in ectopic ventral or posterior locations in relation to the DZ protoglomerulus than in uninjected controls and *ast*-/-. **(C, D)** Misprojections of DZ-projecting BacOR130-1 labeled axons in *robo1* F0 KD; ast-/- embryos and *ast*-/- were more likely to be classified as direct errors instead of branching or escapes from the DZ protoglomerulus. There were mild branching and DZ protoglomerular escape phenotypes in *robo1* F0 KD; *ast*-/- embryos. Statistical comparisons were performed using a two-tailed Fisher’s Exact test with * denoting p ≤ 0.05, ** denoting p ≤ 0.01, *** denoting p ≤ 0.001, and **** denoting p ≤ 0.0001.

**Fig S3.**
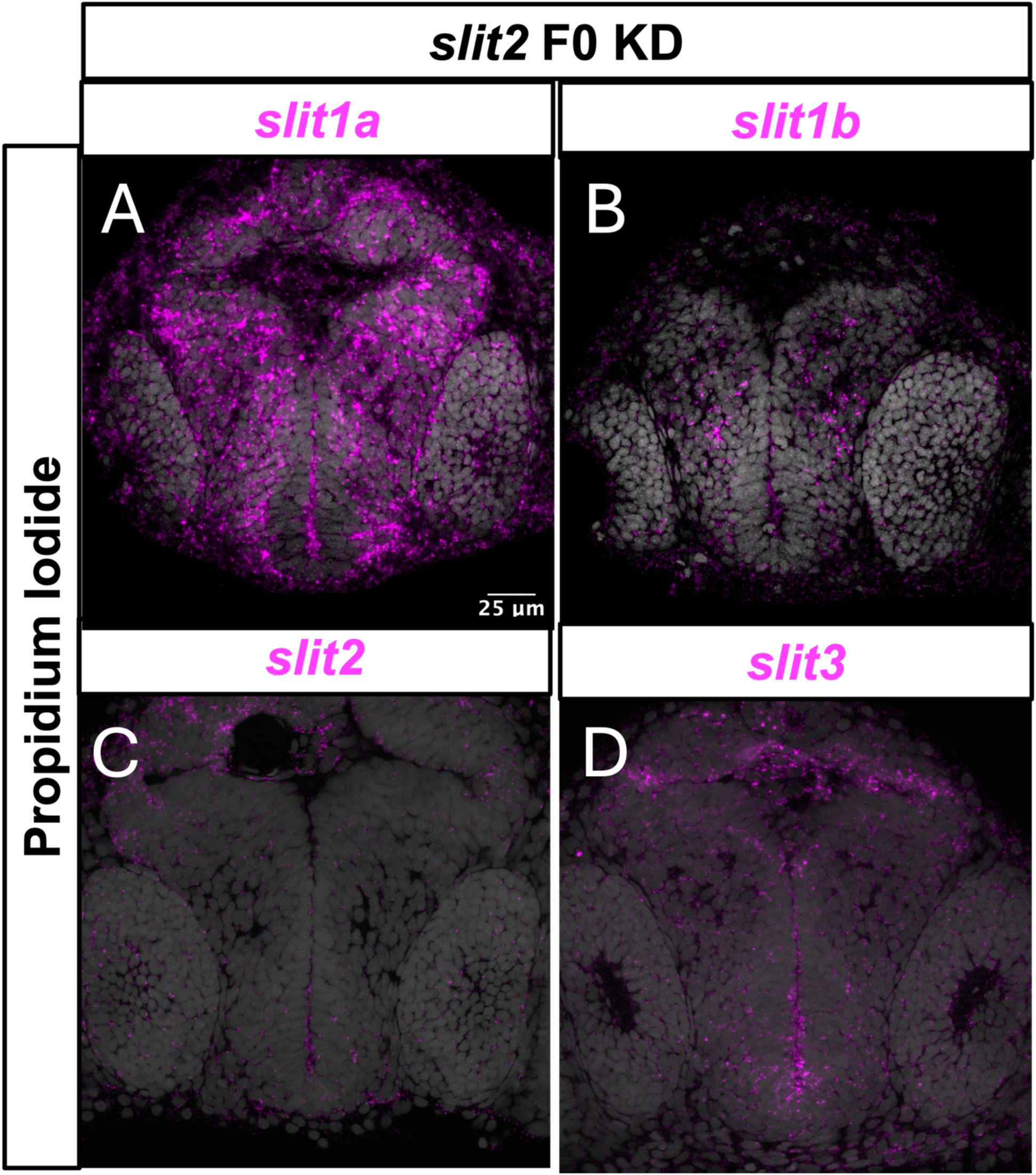
S*lit1a* and *slit3* expression are upregulated in *slit2* F0 KD embryos. Fluorescent *in situ* hybridization was performed at 48 hpf in *slit2* F0 knockdown embryos. Hybridization was performed at the same time in parallel as the preparations presented in Figure 7. Representative 3-µm optical maximum-intensity projections from individual preparations are shown. Upregulation of *slit1a* and *slit3* mRNA in *slit2* knockdowns was observed in 3 independent experiments.

**Fig S4.**
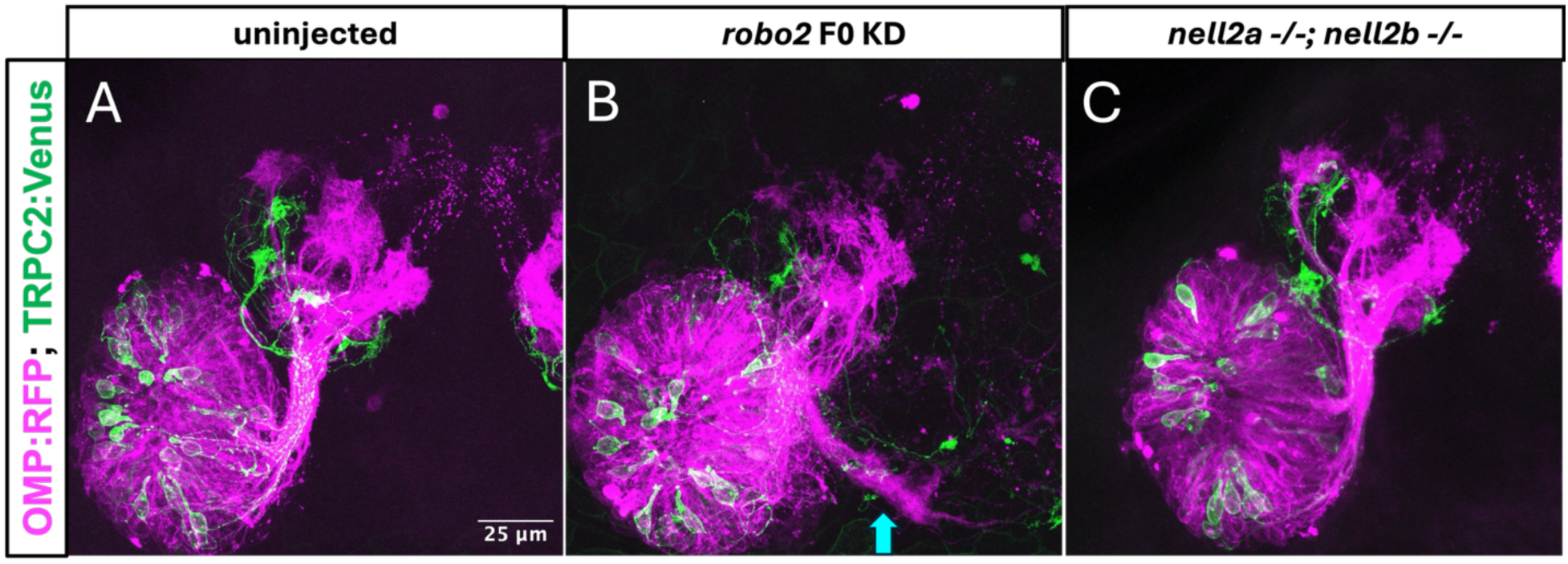
*Nell2a-/-; nell2b-/-* does not phenocopy *robo2* F0 KD. **(A)** Representative 3D projection of an optical z-stack of the zebrafish wild-type olfactory system at 72 hpf. **(B)** The protoglomeruli appear disorganized and a subset of labelled OSN axons misroute posteriorly in *robo2* F0 knockdowns fish. Ventral and posterior axon misprojections are observed in *robo2* F0 knockdowns and is indicated by the cyan arrow. **(C)** The protoglomeruli are not disorganized in *nell2a-/-;nell2b-/-* fish and no ventral and posterior axon misprojections are observed.

